# Characterising modulatory effects of transcranial random noise stimulation using the perceptual template model

**DOI:** 10.1101/2022.09.27.509805

**Authors:** Stephanie Gotsis, Jeroen van Boxtel, Christoph Teufel, Mark Edwards, Bruce Christensen

## Abstract

Neural noise is an inherent property of all nervous systems. However, the mechanisms by which such random and fluctuating neural activity influences perception are still unclear. To elucidate the relationship between neural noise and perceptual performance we require techniques that can safely manipulate neural noise in humans. Transcranial random noise stimulation (tRNS), a form of electrical brain stimulation, has been proposed to increase neural noise according to principles of stochastic resonance (SR); where small to moderate intensities of tRNS improve performance, while higher intensities are detrimental. To date, high intensity (i.e., >2mA) tRNS effects on neural noise levels have not been directly quantified, nor have the detrimental effects proposed by SR been demonstrated in early visual processing. For this purpose, we applied a maximum current intensity of 3mA high-frequency tRNS to the visual cortex (V1) during an orientation discrimination task across increasing external visual noise levels, and fit the perceptual template model to contrast thresholds to quantify intrinsic mechanisms related to noise underlying changes in perceptual performance. We found that tRNS generally worsened perceptual performance by increasing observer’s internal noise and reducing the ability to filter external noise compared to sham. While most observers experienced detrimental effects, others demonstrated improved perceptual performance (i.e., reduced internal noise and better noise filtering). Preliminary evidence suggests that individual baseline internal noise levels may drive the observed beneficial or detrimental observer responses to tRNS. These findings have important implications for the application of tRNS to investigate the impact of internal noise and noise filtering processes on perception.

## Introduction

Neural noise is commonly described in terms of stochastic variability in neuronal firing, where systems with more noise will have greater variation in neuronal activity [1, 2]. An ongoing debate in human perception is how neural noise affects mechanisms involved in perception as causal links have not been clearly established. In the past, research has primarily relied on clinical observations to understand neural noise effects on perception, where disorders such as schizophrenia [3], Parkinson’s disease [4], migraine [5], and dyslexia [6], characteristically present with ‘noisy’ neural processing and disturbed perceptual experiences. However, there are many comorbidities in these conditions like general cognitive and performance deficits [7] that may impact our ability to accurately measure this relationship. Accordingly, we need to establish experimental techniques that can safely increase neural noise in the healthy brain to explore its effects on perceptual performance more directly.

Transcranial random noise stimulation (tRNS) is a non-invasive electrical brain stimulation method that has been shown to impact perceptual performance across several areas, including facial identification, perceptual learning, and perceptual decision-making [8–10]. While the specific mechanisms by which tRNS modulates brain function are yet to be established (*see supplementary information 1*), previous studies suggest that the perceptual effects of tRNS manifest via the modulation of internal neural noise according to the principles of stochastic resonance (SR; [11, 12]). SR leads to a non-monotonic relationship between noise and perceptual performance in the shape of an inverted-U function [13]. The non-monotonic dependence occurs for sub-threshold signals, when the addition of a certain amount of neural noise pushes a weak signal above the threshold for excitation (i.e., quiescent sensory neurons would now have sufficient power to generate an action potential) [14]. For lower levels of noise the signal (plus noise) would remain sub-threshold, while higher levels of noise would obscure the signal.

Van der Groen and Wenderoth [15] demonstrated tRNS effects resembling SR by applying high-frequency tRNS (101 to 640Hz; hf-tRNS) to V1 in human participants during a visual detection task. In this study moderate intensities of hf-tRNS (0.5mA and 1mA) lead to improved performance, while a larger hf-tRNS intensity (1.5mA) lead to perceptual performance comparable to baseline. These changes to perceptual performance are consistent with SR and can be attributed to the hf-tRNS effects on the primary visual cortex. However, SR not only predicts increased performance for moderate noise levels but also decreased performance for higher noise – which was not demonstrated here. Moreover, no direct or computational measure was acquired to quantify neural noise levels in this study.

In this regard, the perceptual template model (PTM) has been used to quantitatively estimate the intrinsic properties of observers under sham (0mA) and active (2mA) hf-tRNS applied to V1 during completion of a visual orientation discrimination task [16]. The PTM assumes that visual processing systems contain a constant amount of internal noise, and that perceptual performance is determined by the type of noise (i.e., internal or external) that dominates the system [17]. To illustrate this relationship, Figure 1 plots contrast thresholds across increasing external visual noise, in a so-called threshold versus external noise contrast (TvN) function [17]. The ‘elbow’ or inflection point of the curve in such functions, where external noise begins to detrimentally affect perceptual performance, is utilised to mathematically estimate the magnitude of *equivalent* internal noise in the system [17]. The PTM suggests that differences in the level of equivalent internal noise and the shape of the TvN function stem from three sources: internal additive noise, internal multiplicative noise, and external noise filtering. Internal additive noise most closely aligns with neural noise as typically discussed in the literature, reflecting noise in the system that is independent of the input and that underlies the stochastic and variable nature of internal responses. By contrast, internal multiplicative noise is dependent on the input and is analogous to contrast gain control mechanisms affecting the system’s responsiveness to stimulus contrast. Lastly, the PTM also considers the impact of external noise filtering, where the addition of external noise may be counteracted by the system’s inherent ability to separate relevant from irrelevant sensory information [17].

**Figure 1.**
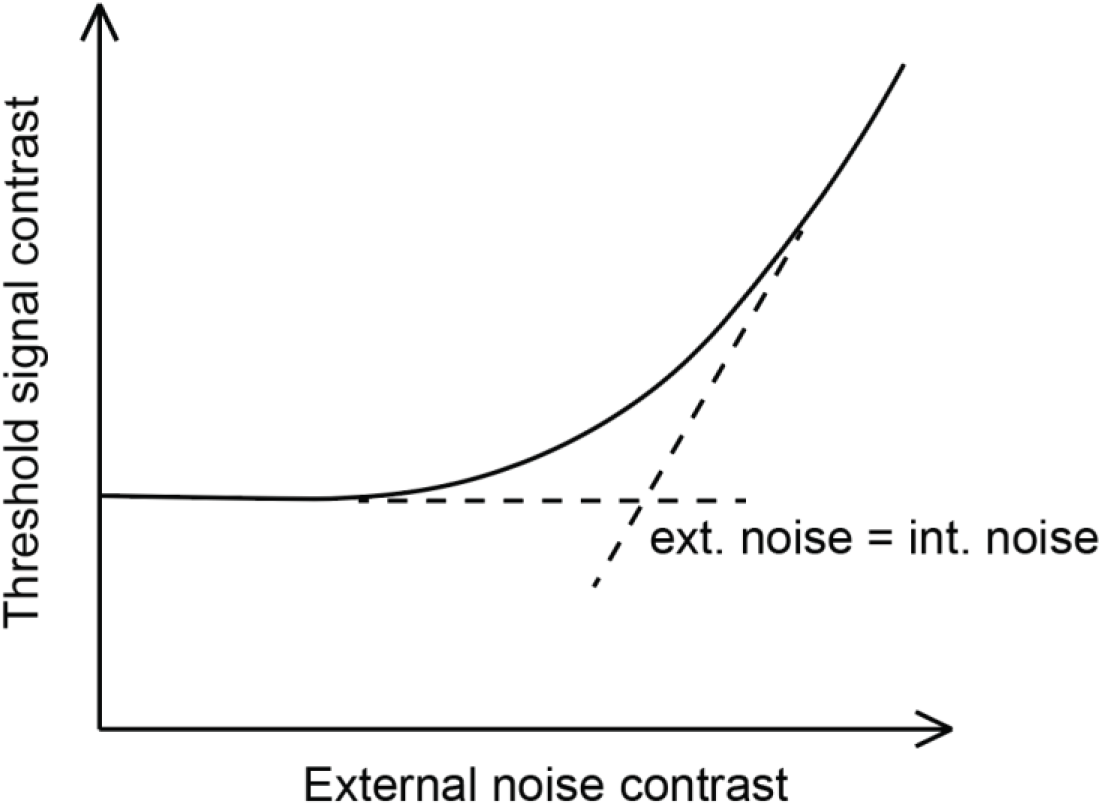
Threshold versus noise contrast (TvN) function. The threshold signal contrast reflects the amount of contrast required to maintain a given performance level (accuracy) as external noise is added to the stimulus. The TvN function is initially characterised by a flat slope in the presence of minimal external noise. This is explained by the fact that performance is dominated by (largely constant) internal noise in the system. The addition of external noise has little to no effect on perceptual performance so long as it does not exceed the internal noise level. As the external noise exceeds the internal noise, it impedes perceptual performance, leading to an increasing slope or ‘arm’ of the curve. The inflection point indicates the point at which internal neural noise is equivalent to the external noise added.

Using the PTM, Melnick et al. [16] did *not* observe significant effects of 2mA hf-tRNS on internal additive noise, with no difference in performance noted across low levels of external noise between stimulation conditions (Figure 1). One possible explanation for this lack of an effect is that the used current intensity was not large enough to increase internal noise to a level that detrimentally affects perceptual performance. This explanation is consistent with findings from van der Groen and Wenderoth [15], who found no effects of performance at 1.5mA. Interestingly, however, Melnick et al. [16] did find that 2mA hf-tRNS improved performance under high external noise levels compared to sham. This finding is consistent with an increased ability to filter external noise. Melnick et al. [16] propose that if hf-tRNS modulates neural activity in a way that is similar to external visual noise the system may be subjected to neural adaptation when concurrently presented. Specifically, Melnick’s [16] presentation of *continuous* hf-tRNS to the cortex over an extended period (~20 minutes) might have minimised neuronal responsiveness in V1 to the external visual noise presented, allowing for better detection of the signal at higher external noise levels. Hf-tRNS applied intermittently may reduce or prevent this from occurring [15].

The current literature provide some evidence to suggest that hf-tRNS current intensity may affect perceptual performance by increasing internal neural noise according to the principles of SR. However, a key aspect in SR – that high levels hf-tRNS induce high amounts of internal noise that can detrimentally affect perception – has not been shown in contrast detection tasks. We therefore aimed to investigate if hf-tRNS could be employed to detrimentally impact perceptual performance by applying a maximum current intensity of 3mA hf-tRNS intermittently to the V1 of healthy observers. To better understand the mechanisms underlying any change in performance, we computationally quantified internal additive noise, internal multiplicative noise, and external noise filtering levels using the PTM. We hypothesised that intermittent 3mA hf-tRNS would detrimentally affect perceptual performance by increasing internal additive noise and reducing participant’s ability to filter external noise. The current theoretical framework does not allow to make specific predictions with respect to the possible effects of hf-tRNS on multiplicative noise.

## Methods

### Participants

Forty-one healthy observers with normal or corrected-to-normal vision served as participants. All participants met the standard safety-related inclusion criteria for tRNS and were exposed to both sham (0mA) and active (3mA) tRNS conditions. The average participation duration was two-hours, and participants received university course credits or payment as remuneration. This study was approved by the Australian National University Human Research Ethics Committee, Canberra Australia (2018/559), and informed written consent was obtained from all participants prior to commencing the experiment. Participants were aged between 18 to 33 years old (*M* = 21.46, *SD* = 3.71), and were primarily female (~73%).

### Apparatus and Stimuli

All computer-based tasks took place in a quiet and dark room. Visual stimuli were generated using Matlab version R2013b and the Psychtoolbox extensions [18–20]. Stimuli were presented on a Compaq colour monitor (P1220) with a calibrated linearized output at a resolution of 1280 x 1024 pixels, and a refresh rate of 85Hz. Contrast was manipulated using the Bits# Stimulus Processor (Cambridge Research Systems). Luminance calibration was performed using a psychophysical procedure in combination with a photometer [21]. A chin rest fixed the viewing distance at 60 cm, with each pixel subtending 0.028 deg. The signal stimulus was a Gabor grating with a cardinal (vertical or horizontal; *θ* = 0 deg or 90 deg) orientation presented in the centre of the screen with a background luminance display (*l*0) was 25 cd/m^2^, Gabor spatial frequency *f* = 4 c/deg, and a Gaussian envelope of *σ* = 0.25 deg. The contrast was determined for every trial by the Psi adaptive staircase procedure (see below).

The signal Gabor patch was temporally sandwiched between two independent external noise samples (Figure 2A). The noise samples had an identical Gaussian envelope to the signal Gabor patch and were constructed using 4×4 pixel elements and sampled from a Gaussian distribution with a mean of 0 and standard deviation of 0, 0.02, 0.04, 0.08, 0.12, 0.16, 0.25, and 0.33. The maximum standard deviation of the external noise was 0.33 to conform to a Gaussian distribution.

**Figure 2.**
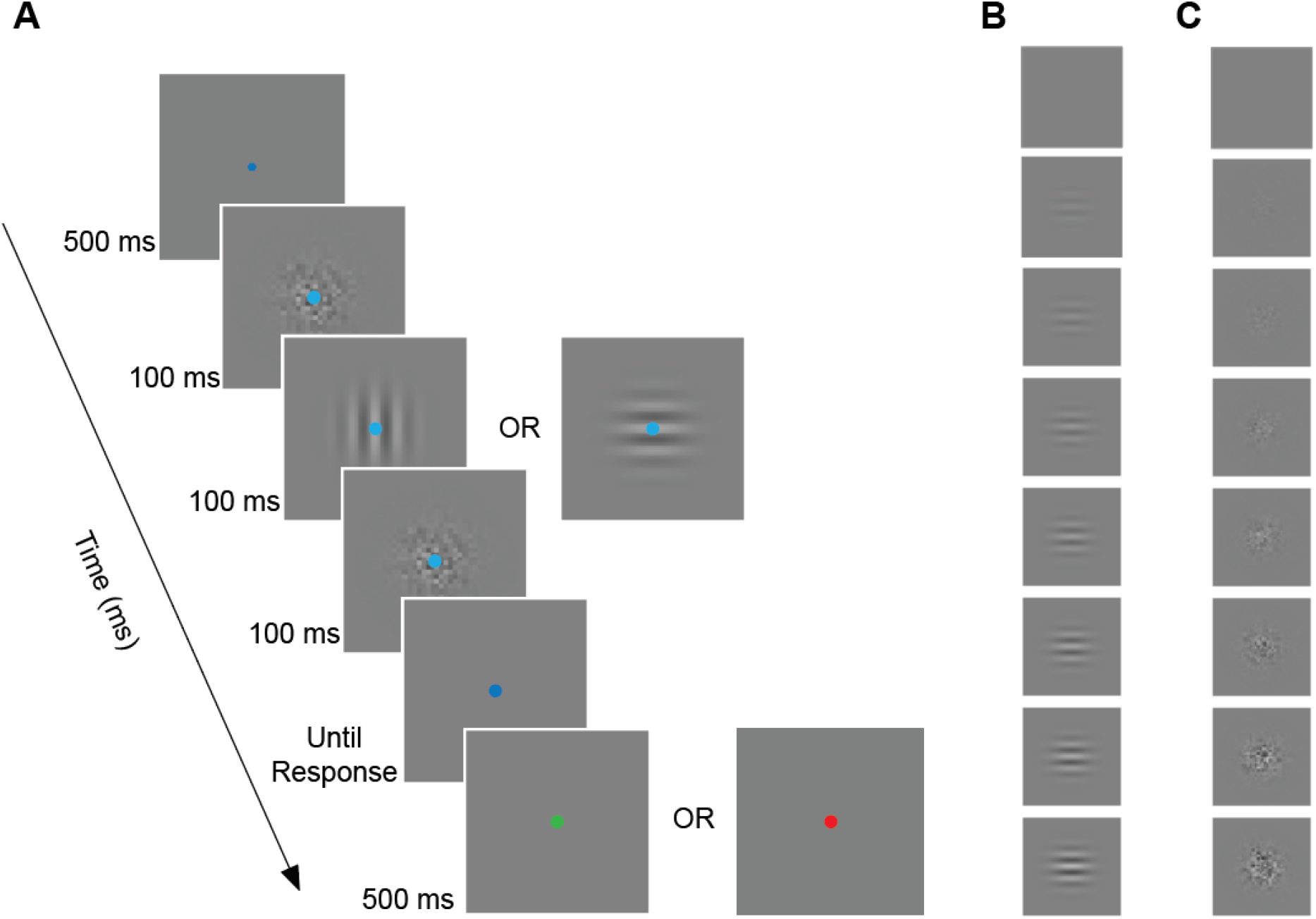
Psychophysical task and features. (A) Each trial began with a central dark-blue fixation dot. The stimulus sequence coincided with a fixation colour change to light-blue, and comprised a series of three frames: Gaussian pixel noise, oriented Gabor grating (vertical or horizontal), Gaussian pixel noise. For each trial, the subjects’ task was to judge whether the grating was vertically or horizontally oriented. Feedback was then provided in the form of a fixation colour change to green or red, indicating correct or incorrect responses. Given the short duration of each frame in the stimulus sequence, the perceived stimulus appeared as a single image (i.e., Gabor + noise). From top to bottom: (B) a vertically oriented signal Gabor patch with increasing contrast; (C) Eight external noise images with increasing external noise (standard deviation: 0%, 2%, 4%, 8%, 12%, 16%, 25%, and 33%).

### Procedure and design

This study was comprised of a small practice session, and two experimental sessions (sham-tRNS and active-tRNS sessions, counterbalanced). A two-alternative forced choice (2-AFC) task required participants to judge if the Gabor stimulus had a vertical or horizontal orientation (Figure 2A). Participants made their response by pressing keys 1 or 2 on the number keypad for vertical or horizontal orientations, respectively. Each trial took ~2 seconds, including the participant response time (typically less than 1 s).

The practice session introduced the 2-AFC task (20 trials) that would then be used in the experimental sessions (Figure 2A). Sham-tRNS and active-tRNS sessions applied the Psi adaptive staircase method (Palamedes toolbox; [22]) to adjust the stimulus contrast on every trial (Figure2B), and estimate the contrast thresholds (i.e., the minimal signal strength required for the subject to accurately detect the orientation of the Gabor stimulus) for each external noise condition (Figure 2C). Psychometric functions for eight external noise conditions were estimated, using an interleaved design of 60 trials per noise condition. In total, 480 trials over 10 blocks containing 48 trials were completed. Between each block, participants were required to take a minimum 60 second break before proceeding. Weibull functions were used to estimate contrast thresholds for three different performance criterion levels (*d’·.* 0.78, 1.35, and 2.07), corresponding to 65%, 75% and 85% performance accuracy:

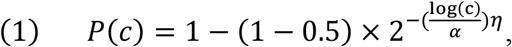

where the *P* is the percent correct, *c* is the stimulus contrast, *α* is the threshold performance, and *η* is the slope of the psychometric function in equation (1).

### tRNS protocol

Participants received an active-tRNS current (3mA) during one of the experimental sessions (active session), and sham-tRNS (0mA) in the other (sham session). For both sessions, two 5×7 cm electrodes were positioned on the participants scalp and held in place with a soft rubber headband. The electrodes were coated with an electrode gel (Signa gel; Parker Laboratories Inc) to reduce skin impedance. Based on the 10-20 EEG system, the anode was positioned at the occipital region Oz and the cathode at the vertex of the scalp (Cz). This setup is an established method for stimulation of the primary visual cortex [15, 23].

During the active session, 3mA of high-frequency tRNS (hf-tRNS; 101-640Hz) was administered to participants during the task and terminated during breaks between blocks. Investigations applying such *intermittent* stimulation could be used to minimise the possibility of adaptive mechanisms taking effect [15]. Stimulation was triggered by a spacebar press required to start each block and ended at the end of each block or after a maximum of 105 seconds (including ramp-up stimulation phase of 5 seconds). The impedance of each electrode was checked before and during stimulation to ensure it was under 10kΩ. The stimulation equipment set-up for sham-sessions was identical to the active-session, however no current (0mA) was presented.

### Perceptual Template Model Analysis

The effects of stimulation on perceptual performance were estimated by fitting individual data with the PTM and comparing active-tRNS versus sham-tRNS. The PTM model quantifies the extent to which perceptual performance under each condition was limited by internal additive noise, internal multiplicative noise, or external noise filtering. Lu and Dosher [17] therefore characterise perceptual thresholds (c_τ_) by equation (2):

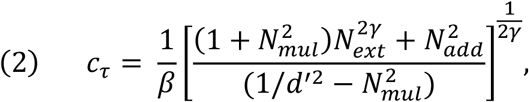

where an input consisting of signal and external noise (*N_ext_*) is received by an observer and transformed into an internal representation. This representation is passed through a perceptual ‘template’, which can be understood as a processing filter with selectivity for certain visual characteristics (e.g., orientation, colour, and spatial frequency). The system’s ability to filter external noise (*N_ext_*) will affect its ability to match the signal with the template. The signal can be enhanced by a gain factor *β* depending on how well the input is matched to the template. This template output will undergo another transformation through a nonlinear transducer function *γ*, accounting for the nonlinear properties of the visual system. Two sources of internal noise, that is additive (*N_add_*) and multiplicative (*N_mul_*), are added to the transformed signal.

TRNS effects were characterised by introducing three coefficient indices (*Aa(stim), Am(stim), and Af(stim)*) to the conventional equation (2). As seen in equation (3), each coefficient is multiplied by the corresponding source of noise: additive noise (*N_add_*), multiplicative noise (*N_mul_*), or external noise filtering (*N_ext_*):

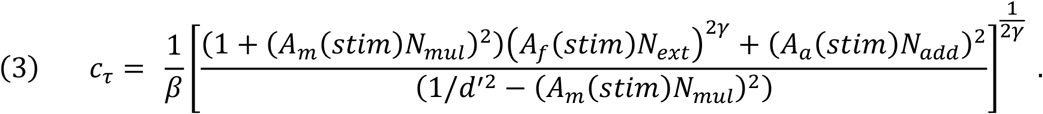

The influence of these noise types on performance between sham and active-tRNS conditions was determined by fixing sham stimulation coefficient indices (*Aa(sham), Am(sham)*, and *Af(sham)*) to 1, while the indices for active stimulation (*Aa(active), Am(active),* and *Af(active)*) were free to vary. The resulting coefficients, therefore, describe the relative difference in the effects of tRNS between the stimulation groups; i.e., parameters greater than 1 suggest that the active stimulation produced higher amounts of the respective noise types compared to the sham stimulation. Eight forms of the PTM were considered on the basis that the coefficient indices for active stimulation could vary or be fixed to 1. For instance, a null model assumes that there are no group differences between stimulation conditions and, therefore, four free parameters (*N_mul_, N_add_, β,* and *γ*). Alternatively, the fullest model has up to seven free parameters.

The weight of each threshold data point fit by the PTM varied according to the goodness of fit of the Weibull function used to obtain the threshold estimate. Weibull function fits that were <0.001 were set to 0.001 as a minimum, and weights were then normalised relative to the maximum weight. Thus, threshold estimates with better fitting Weibull functions were weighted more heavily, compared to those obtained from poorer fitting functions. This ensured that the most reliable threshold estimates had the most influence in the model.

A least square procedure was used to fit the PTM to the TvN data for each participant. The least square difference between the log of the measured threshold contrast and the log of the model-predicted threshold contrast was minimised to find the best fit for the reduced and full models. The *r^2^* statistic was used to measure the goodness-of-fit using equation (4):

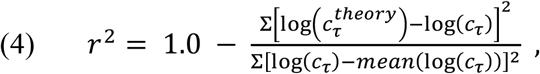

where, Σ and *mean()* were applied across all external noise levels, stimulation conditions, and performance levels.

The best-fitting PTM was determined across participants by comparing the Akaike Information Criterion (AIC; [24]) for all eight PTM variations. The AIC for each model was calculated for individual participants and averaged. The model with the a) the lowest AIC estimate, which b) meaningfully deviated from the other models (i.e., a difference greater than 2), was identified as having the best fit (see Wagenmakers and Farrell [24] for details).

## Results

Upon inspection of the PTM results for each participant (*N* = 41), we identified 6 participants whose data could not be fit by the PTM to a moderate or high standard (i.e., *r*^2^ <.5) [25]. As a result, these datasets were excluded from analyses, leaving 35 participants in the final analysis.

AIC model selection was applied to identify which of the eight PTM variations best accounted for differences between stimulation conditions. The PTM variation including internal additive noise and external noise filtering only was identified as the best model (i.e., model AF; Table 1). AIC model selection performed on individual participants also supported this finding (*see supplementary information 2).* This suggests that 3mA hf-tRNS meaningfully differed from sham-tRNS in levels of internal additive noise and external noise filtering produced, but not in the amount of internal multiplicative noise produced. An assessment of AIC weights (w(AIC); Table 1) further supports this conclusion, where model AF, accounted for approximately 31% of the total explanation that can be found in the full set of models.

**Table 1.** Akaike Information Criterion (AIC) estimates for PTM variations.

|  | Model Variations |  |  |  |  |  |  |  |
| --- | --- | --- | --- | --- | --- | --- | --- | --- |
|  | AFM | AF | AM | FM | A | F | M | Null |
| AIC | -38.19 | -40.17 | -35.99 | -38.17 | -36.80 | -39.90 | -36.00 | -36.55 |
| $\Delta\text{AIC}$ | 1.98 | 0.00 | 4.17 | 1.99 | 3.37 | 0.27 | 4.17 | 3.62 |
| $w(\text{AIC})$ | 0.12 | 0.31 | 0.04 | 0.11 | 0.06 | 0.27 | 0.04 | 0.05 |

*Note.* The best fitting model has the smallest AIC estimate that meaningfully deviates from other models by a factor of $\sim 2$ or more ( $\Delta\text{AIC}$ is the difference between the best model and each other model); $w(\text{AIC}) = \text{AIC}$ weight is proportional to the total amount of predictive power provided by the full set of models contained in the model being assessed [24]. AFM – fullest model with Aa, Af, and Am coefficients free to vary; AF – Aa and Af coefficients free to vary; AM – Aa and Am coefficients free to vary; FM – Af and Am coefficients free to vary; A – Aa coefficient free to vary; F – Af coefficient free to vary; M – Am coefficient free to vary; Null – Aa, Af, and Am fixed to 1.

Further investigations of the best fitting PTM demonstrated characteristic nonlinear TvN functions, where contrast thresholds increased as a function of increasing levels of external noise contrast (Figure 3; a representative subject). Internal additive noise estimates showed an average 19% increase under 3mA hf-tRNS conditions compared to sham-tRNS. This effect is demonstrated in the TvN function by the increase in contrast threshold for 3mA hf-tRNS across low external noise levels compared to sham-tRNS (Figure 3). External noise filtering estimates showed a considerably smaller average 3% increase under 3mA hf-tRNS conditions compared to sham-tRNS. This small difference was demonstrated as a slight increase in contrast thresholds across high external noise conditions in the TvN functions (Figure 3).

**Figure 3.**
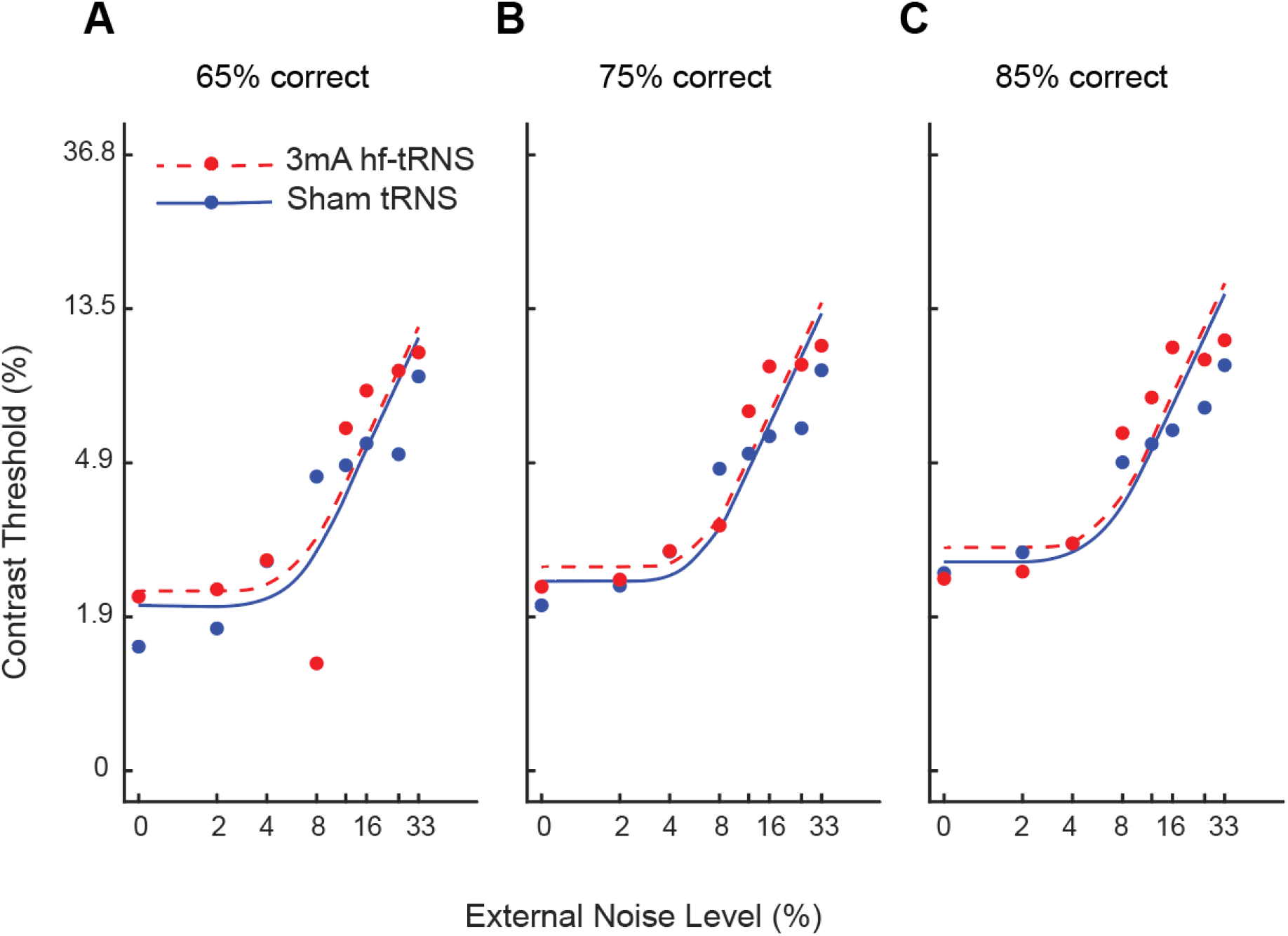
Contrast threshold versus external noise contrast (TvN) functions. TvC functions at (A) 65%, (B) 75%, and (C) 85% performance accuracy. This representative participant reflects the general findings, displaying ~20% increase in additive noise and ~3% increase in external noise filtering under 3mA hf-tRNS (red circle + dashed line) compared to sham (blue circle + solid line) estimated by the PTM. Eight external noise levels (standard deviation: 0%, 2%, 4%, 8%, 12%, 16%, 25%, and 33%)

PTM estimates across individual participants showed considerable variability (Figure 4A and 4B). Overall, 3mA hf-tRNS appeared to increase internal additive noise and external noise filtering estimates in ~55% of participants (Figure 4A and Figure 4B, respectively).

**Figure 4.**
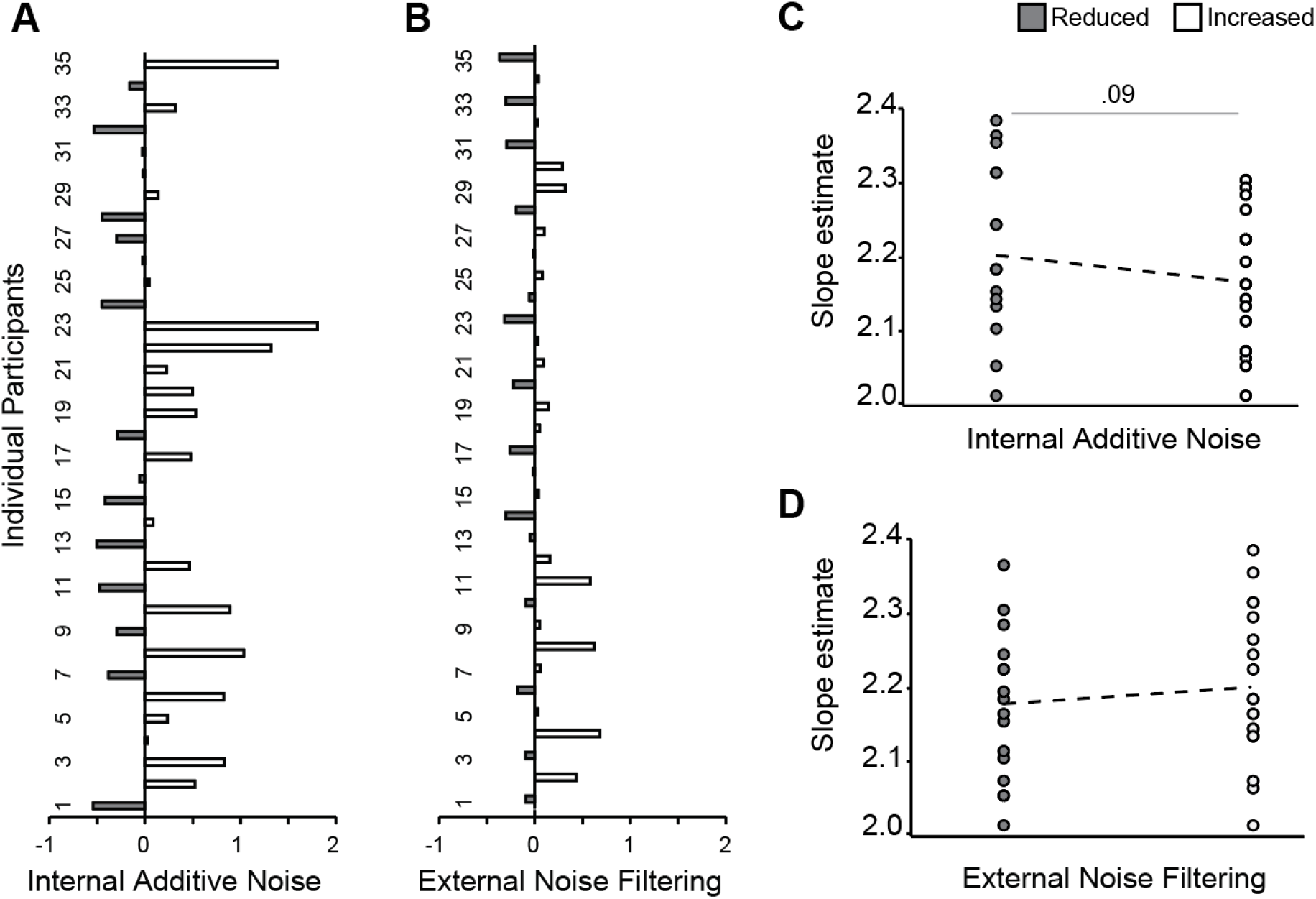
Individual variability of tRNS effects. (A) PTM internal additive noise and (B) external noise filtering estimates after 3mA hf-tRNS compared to sham-tRNS. A negative shift from zero is indicative of a reduction (n=16), while a positive shift is indicative of an increase (n=19) in the corresponding noise type. Please note that an increase in external noise filtering is indicative of poorer filtering. (C) and (D) Baseline internal noise is represented as the slope estimate of the psychometric function for the zero external noise condition under sham hf-tRNS for all participants. We present the correlation between slope estimates and reduced or increased internal additive noise (C) or external noise filtering (D) PTM estimates.

The spread of the psychometric function was used to explore whether baseline internal noise levels were associated with reduced (i.e., beneficial) or increased (i.e., detrimental) PTM estimates. The spread of the psychometric function, which is the slope, for the threshold obtained under sham conditions with zero external noise was used as a measure of baseline internal noise ([13]; *see supplementary information 3).* A larger spread (i.e., flatter/smaller slope) would be indicative or more variance, and thus greater internal noise. Point biserial correlations showed that smaller slope estimates, indicative of greater baseline internal noise, may relate to increased internal additive noise estimates produced by the PTM under 3mA hf-tRNS (*r* = -.228, *p* = .094; Figure 4C). With regards to external noise filtering, a bivariate correlation showed a weak, positive, non-significant correlation between the slope estimate with external noise filtering outcomes (*r* = .148, *p* = .198; Figure 4D).

Other participant characteristics including age, gender, intelligence, and personality traits were also explored with bivariate correlations. All correlations were non-significant (*see supplementary information 4*).

## Discussion

The present study provides evidence that high intensity (3mA) hf-tRNS over V1 can worsen perceptual performance across increasing external noise levels in a contrast detection task. The PTM revealed that detrimental effects of tRNS on performance resulted from increased internal additive noise and reduced ability to filter external noise. No effects of 3mA hf-tRNS were seen on internal multiplicative noise. These findings support van der Groen and Wenderoth [15] work suggesting that hf-tRNS increases neural noise to affect perceptual performance in a way that is consistent with principles of SR. In addition, our results showing poorer external noise filtering allow us to further explore ideas pertaining to hf-tRNS effects on neural adaptation as proposed by Melnick and colleagues. Further investigations of individual participants subsequently offer some preliminary evidence suggesting that individual responses to hf-tRNS may be driven by the baseline noise levels pre-existing in the observers system. These outcomes provide important implications for the application of hf-tRNS as a means of investigating the impacts of noise on perception.

When considered in the context of pre-existing work, our result that 3mA hf-tRNS can be detrimental to performance by increasing internal additive noise provides further evidence for SR as a plausible mechanism underlying hf-tRNS effects on performance. Specifically, previous research by van der Groen and Wenderoth [15] and Melnick [16] were not able to demonstrate the full SR effect showing diminished perceptual performance below that of baseline performance levels under 1.5mA and 2mA hf-tRNS (respectively). Additionally, Melnick et al. found no significant changes in internal additive noise estimates produced by the PTM. It is possible that 2mA hf-tRNS does not modulate internal noise to affect performance, however, it is also possible that the increased noise induced by 2mA hf-tRNS may have resulted in performance that was equivalent to baseline performance; landing on the descending arm of the SR curve. As PTM estimates are based on differences in perceptual performance between active and sham conditions, if there is no perceived difference in performance between stimulation groups across low external noise conditions the PTM would subsequently estimate no change to the internal additive noise estimate. Thus, it is possible, if not likely, that internal additive noise did increase under 2mA hf-tRNS; however, this was not visible as a *change* in performance and therefore not detected by the PTM.

The exploration of individual responses to hf-tRNS demonstrated substantial variability across participants, with some participants showing improved perceptual performance that was characterised by the PTM as a reduction in internal additive noise. The PTM assumes optimal criterion setting and that increased internal noise will result in detrimental effects to performance. Therefore, it cannot model the improved effects of SR. Approaches similar to the PTM, such as the linear amplifier model, have been shown to misestimate the level of internal noise, when conditions allow for SR [26]. This has serious implications when interpreting perceptual performance that results in a reduction of the estimated internal additive noise parameter. In the case of SR, improved performance might not necessarily be attributed to reduced internal noise in the system, but rather, low to moderate *increases* in internal noise. Thus, if hf-tRNS increases internal noise leading to SR, and improves perceptual performance, this would be inaccurately represented as a reduction to internal additive noise estimated by the PTM. Based on the reduced internal additive noise estimate alone, one cannot determine if the reduced additive noise estimate simply represents improved performance resulting from *reduced* neural noise, or if it is a representation of improved performance resulting from *increased* neural noise according to SR. If the latter is true, the average increase in additive noise reported in our study would be largely underestimated, as the reduced additive noise estimates produced by the PTM may actually reflect increased noise resulting in improved performance in a SR model.

The results also indicate that exposure to 3mA hf-tRNS generally diminished participant’s ability to filter external noise. This is shown as a slight increase in contrast thresholds (i.e., worse performance) at higher external noise levels under 3mA hf-tRNS compared to sham. This finding is inconsistent with Melnick and colleagues [16] that demonstrated the opposite effect; i.e., improved external noise filtering effects from 2mA hf-tRNS. Melnick proposed that if hf-tRNS and external white noise have similar mechanisms of action to affect perceptual performance (i.e., they both increase random and fluctuating neural activity in the cortex), they may be susceptible to adaptation when presented concurrently. This idea is potentially supported by van der Groean and Wenderoth [15] that showed increasing current intensities of hf-tRNS affected behavioural performance just as increasing intensities of external white noise (i.e., according to SR). Accordingly, prolonged and continuous presentation of hf-tRNS may affect an observer’s ability to filter external noise through adaptation by minimising neuronal responsiveness to external white noise presented in the visual stimuli [27]. If adaptation is responsible for improved external noise filtering ability shown by Melnick et al. [16], the opposing effect of poorer external noise filtering ability observed in our present study may be due to our stimulation protocol preventing adaptation. Specifically, Melnick [16] presented a persistent current over a 20-minute period during task completion. However, we presented 20 minutes stimulation intermittently, using 2-minute stimulation periods with 1-minute breaks where stimulation ceased. Since a lack of change in the adaptor is a crucial feature for adaptation to occur, the stimulation protocol applied by Melnick and colleagues may have provided a consistent and prolonged ‘noisy’ cortical environment allowing for adaptation to the higher external visual noise levels, and consequently, better external noise filtering. Alternatively, the inconsistent and changing environment produced by the intermittent stimulation presented in our study may have substantially reduced the possibility for any adaption to occur, diminishing the observers’ ability to filter external noise. Unfortunately, our study was not designed to explore the impact of adaptation, as we inter-mixed external noise conditions across task blocks. This would additionally minimise any adaptive mechanisms to the external noise presented, and therefore the split-half analysis that we would apply to explore adaptation across a testing block would not be appropriate. Future research may want to consider this in their experimental design to explore adaptive mechanisms.

Our subsequent investigations of differential hf-tRNS effects on perceptual performance across observers showed that individual baseline internal noise levels were related to hf-tRNS benefits on perceptual performance. Specifically, we found a trend-level relationship where participants with low baseline internal noise levels were associated with improved perceptual performance (i.e., reduced additive noise estimates) under 3mA hf-tRNS compared to sham. In accordance with SR, the addition of hf-tRNS to systems with low baseline noise may have produced a moderate increase in neural noise (quantified as a reduction in additive noise by the PTM) to improve perceptual performance. However, for those with higher pre-existing baseline noise levels in their system, the addition hf-tRNS may have increased neural noise beyond the optimal level for performance improvement to detrimental levels. While 3mA hf-tRNS also appeared to have varying effects across observers’ ability to filter external noise, individual baseline noise levels did not appear to explain this observation. Our relatively small sample size did not allow us to further utilise recorded demographic information to explore other individual factors that may contribute to the differential effects of hf-tRNS shown. Future research with larger sample sizes and experimental designs optimised to measure adaptation are recommended to explore these possibilities.

In conclusion, this study shows that visual performance can be detrimentally impacted by 3mA hf-tRNS compared to sham. Modelling with the PTM suggests that the mechanisms driving this effect are increased internal additive noise and impaired external noise filtering. In the context of previous work, these results further support the notion that hf-tRNS increases additive noise according to current intensity to affect performance in line with SR. However, given that the PTM does not account for SR, our findings may be an underestimate of the full effect. We additionally provide some preliminary evidence suggesting that pre-existing baseline noise levels of observers may drive whether observers experience beneficial or detrimental effects to hf-tRNS. Future work could focus on designing experiments that allow for more in-depth exploration of relationships between baseline noise, SR, and adaptive mechanisms with hf-tRNS.

