## Supplementary information for "Characterising modulatory effects of transcranial random noise stimulation using the perceptual template model"

*1) Transcranial random noise stimulation*

Transcranial random noise stimulation (tRNS) generates random electrical frequencies across specified ranges (e.g., low-frequency, 0.1 to 100Hz; high-frequency, 101 to 640Hz) at current intensities typically between 0 to 3mA. These random electrical frequencies oscillate between two targeted cortical brain regions corresponding to the locations of two electrodes placed on the scalp. An important mechanistic candidate of tRNS is increased neuronal excitability reflecting internal noise; potentially arising from multifaceted neural alterations including the weak neuronal membrane depolarisation, augmentation of voltage-gated sodium channels, broadening of neuronal tuning bandwidths, and/or alteration of neural oscillations (Antal & Herrmann, 2016).

*2) AIC Model Selection for Individual Participants*

Table 1

*Individual Akaike Information Criterion (AIC) estimates for PTM variations.*

|  | Model Variations | | | | | | | |
| --- | --- | --- | --- | --- | --- | --- | --- | --- |
| ID | AFM | AF | AM | FM | A | F | M | NULL |
| 1 | -17.49 | -19.49 | -18.70 | -14.62 | -20.70 | -16.46 | -15.78 | -15.97 |
| 2 | -79.16 | -81.16 | -68.85 | -78.12 | -69.21 | -79.77 | -69.10 | -65.44 |
| 3 | -103.60 | -105.62 | -101.48 | -104.84 | -101.55 | -106.75 | -103.48 | -103.35 |
| 4 | -35.55 | -37.44 | -24.21 | -37.54 | -23.22 | -39.43 | -26.12 | -23.03 |
| 5 | -10.66 | -12.66 | -12.58 | -12.30 | -14.58 | -14.30 | -14.22 | -16.18 |
| 6 | -19.41 | -21.41 | -18.47 | -18.00 | -19.94 | -19.19 | -17.17 | -19.17 |
| 7 | -36.35 | -38.35 | -37.90 | -36.34 | -39.90 | -37.43 | -37.23 | -39.22 |
| 8 | 18.90 | 17.05 | 23.10 | 18.38 | 22.27 | 16.38 | 21.99 | 22.47 |
| 9 | -70.69 | -72.69 | -72.19 | -70.80 | -74.19 | -72.36 | -72.13 | -74.06 |
| 10 | -18.57 | -20.57 | -19.87 | -16.95 | -21.62 | -18.94 | -18.51 | -20.51 |
| 11 | -65.90 | -67.90 | -40.51 | -64.75 | -38.05 | -66.22 | -42.17 | -39.93 |
| 12 | -49.35 | -51.35 | -49.93 | -50.03 | -50.84 | -51.47 | -51.34 | -50.63 |
| 13 | -37.05 | -39.05 | -38.96 | -36.31 | -40.96 | -38.31 | -37.40 | -38.83 |
| 14 | -15.31 | -17.31 | -10.01 | -17.26 | -9.28 | -19.26 | -12.05 | -11.08 |
| 15 | -30.12 | -32.12 | -31.99 | -28.37 | -33.99 | -30.34 | -30.37 | -32.34 |
| 16 | -23.44 | -25.44 | -25.42 | -25.41 | -27.42 | -27.41 | -27.37 | -29.37 |
| 17 | -14.97 | -16.97 | -11.61 | -15.50 | -11.11 | -17.50 | -12.26 | -12.77 |
| 18 | -33.29 | -35.09 | -35.28 | -33.84 | -36.77 | -35.84 | -35.61 | -37.59 |
| 19 | -37.97 | -39.65 | -39.90 | -39.73 | -40.13 | -40.36 | -41.49 | -40.85 |
| 20 | -24.81 | -26.81 | -23.31 | -25.49 | -24.33 | -27.49 | -24.48 | -26.16 |
| 21 | -19.24 | -21.24 | -20.80 | -20.60 | -22.80 | -22.60 | -22.03 | -23.99 |
| 22 | -77.36 | -79.36 | -79.24 | -71.84 | -81.24 | -73.27 | -73.67 | -73.40 |
| 23 | -5.41 | -7.41 | -4.40 | -2.98 | -5.83 | -4.84 | -3.60 | -5.55 |
| 25 | -31.67 | -33.67 | -33.36 | -28.25 | -35.36 | -30.06 | -29.95 | -30.20 |
| 25 | -21.69 | -23.69 | -23.42 | -23.67 | -25.42 | -25.67 | -25.30 | -27.30 |
| 26 | -63.66 | -65.66 | -65.64 | -65.65 | -67.64 | -67.65 | -67.61 | -69.61 |
| 27 | -27.68 | -29.68 | -28.97 | -29.09 | -30.97 | -31.09 | -30.70 | -32.70 |
| 28 | -34.01 | -36.01 | -33.36 | -32.63 | -35.31 | -34.60 | -31.96 | -32.27 |
| 29 | -95.28 | -97.28 | -84.82 | -96.98 | -79.78 | -98.79 | -81.57 | -79.82 |
| 30 | -9.04 | -11.04 | -8.18 | -11.03 | -9.26 | -13.03 | -10.16 | -10.97 |
| 31 | -14.50 | -16.50 | -11.95 | -16.50 | -13.80 | -18.50 | -13.58 | -14.80 |
| 32 | -58.08 | -60.07 | -60.03 | -55.21 | -62.00 | -56.59 | -56.82 | -58.56 |
| 33 | -59.89 | -61.89 | -47.17 | -60.60 | -41.84 | -62.60 | -47.62 | -43.82 |
| 34 | -59.83 | -61.83 | -61.48 | -61.28 | -63.48 | -63.27 | -63.01 | -65.01 |
| 35 | -54.45 | -56.45 | -39.01 | -51.93 | -37.78 | -51.61 | -36.31 | -37.45 |

Table 2

*Average Akaike Information Criterion (AIC) estimates and weight calculation for PTM variations.*

|  | Model Variations | | | | | | | |
| --- | --- | --- | --- | --- | --- | --- | --- | --- |
|  | AFM | AF | AM | FM | A | F | M | Null |
| aveAIC | -43.06 | -45.06 | -34.10 | -41.40 | -34.35 | -42.71 | -33.39 | -33.30 |
| ∆AIC | 2.00 | 0.00 | 10.96 | 3.66 | 10.71 | 2.35 | 11.67 | 11.76 |
| w(AIC) | 0.20 | 0.54 | 0.00 | 0.09 | 0.00 | 0.17 | 0.00 | 0.00 |

*Note.* aveAIC = the average AIC estimate across all participants. ∆AIC is the difference between the best model and each other model; w(AIC) = is proportional to the total amount of predictive power provided by the full set of models contained in the model being assessed (Wagenmakers & Farrell, 2004). Perceptual Template Model variations: AFM – fullest model with Aa, Af, and Am coefficients free to vary; AF – Aa and Af coefficients free to vary; AM – Aa and Am coefficients free to vary; FM – Af and Am coefficients free to vary; A – Aa coefficient free to vary; F – Af coefficient free to vary; M – Am coefficient free to vary; Null – Aa, Af, and Am fixed to 1.

*3) Spread of the psychometric function as proxy for baseline noise levels*

As outlined by Aihara et al. (2010), the spread of the psychometric function corresponds to trial-to-trial variability, and therefore is assumed to reflect the total noise level in the system. Accordingly, under the sham stimulation condition when zero external noise was applied, the spread of the psychometric function may be interpreted to reflect the internal noise level alone.

*4) Bivariate correlations*

Table 1

*Bivariate correlations assessing the relationship between participant characteristics and hf-tRNS effects on internal additive noise and external noise filtering.*

| Variable | Internal additive noise | External noise filtering |
| --- | --- | --- |
| Internal additive noise | - |  |
| External noise filtering | -.04 | - |
| Age | .26 | .01 |
| Gender | .08 | .28 |
| Education | .29 | .02 |
| FISQ | -.25 | .08 |
| PAS | -.20 | .09 |
| SPQ | .10 | .01 |

*Note.* FISQ = Full Scale Intelligence Quotient; PAS = Personality Assessment Screener; SPQ = Schizotypal Personality Questionnaire.
